## Supplemental Table S1 for "Computational design and interpretation of single-RNA translation experiments"

### Supplementary Table 1.

**Table S1.** Codon usage table calculated from the *Homo sapiens* genome. Table is computed using 93,487 CDS (Coding DNA Sequence), that represent a total of 40,662,582 codons, Nakamura, et al., 2000. Reference [21] in main Text.

| | | | |
|---|---|---|---|
| TTT 17.6 | TCT 15.2 | TAT 12.2 | TGT 10.6 |
| TTC 20.3 | TCC 17.7 | TAC 15.3 | TGC 12.6 |
| TTA 7.7 | TCA 12.2 | TAA 1.0 | TGA 1.6 |
| TTG 12.9 | TCG 4.4 | TAG 0.8 | TGG 13.2 |
| CTT 13.2 | CCT 17.5 | CAT 10.9 | CGT 4.5 |
| CTC 19.6 | CCC 19.8 | CAC 15.1 | CGC 10.4 |
| CTA 7.2 | CCA 16.9 | CAA 12.3 | CGA 6.2 |
| CTG 39.6 | CCG 6.9 | CAG 34.2 | CGG 11.4 |
| ATT 16.0 | ACT 13.1 | AAT 17.0 | AGT 12.1 |
| ATC 20.8 | ACC 18.9 | AAC 19.1 | AGC 19.5 |
| ATA 7.5 | ACA 15.1 | AAA 24.4 | AGA 12.2 |
| ATG 22.0 | ACG 6.1 | AAG 31.9 | AGG 12.0 |
| GTT 11.0 | GCT 18.4 | GAT 21.8 | GGT 10.8 |
| GTC 14.5 | GCC 27.7 | GAC 25.1 | GGC 22.2 |
| GTA 7.1 | GCA 15.8 | GAA 29.0 | GGA 16.5 |
| GTG 28.1 | GCG 7.4 | GAG 39.6 | GGG 16.5 |
