## Supplementary figures and images for "Computational design and interpretation of single-RNA translation experiments"

### Supplemental Fig S3

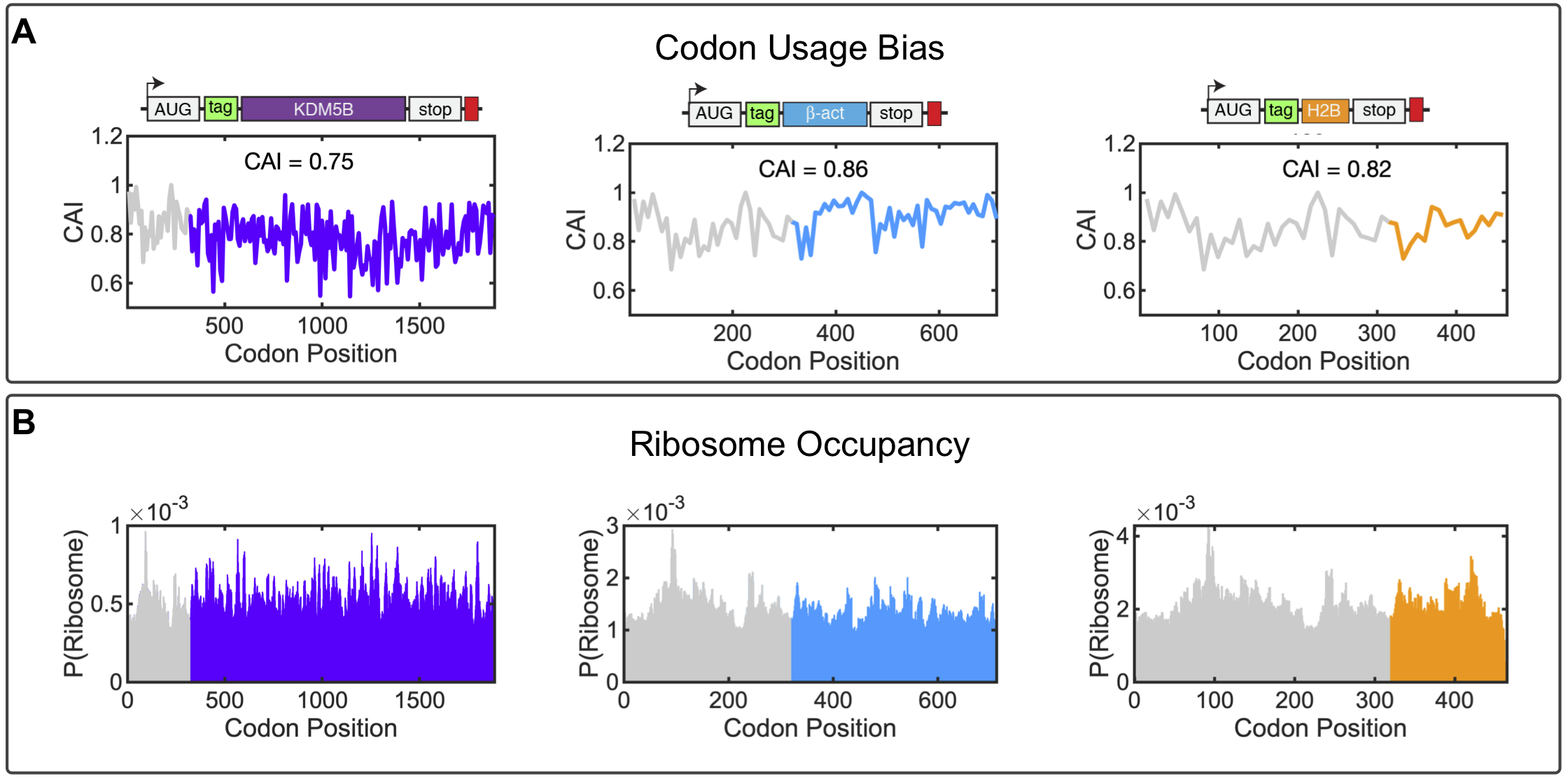

### Supplemental Fig S4

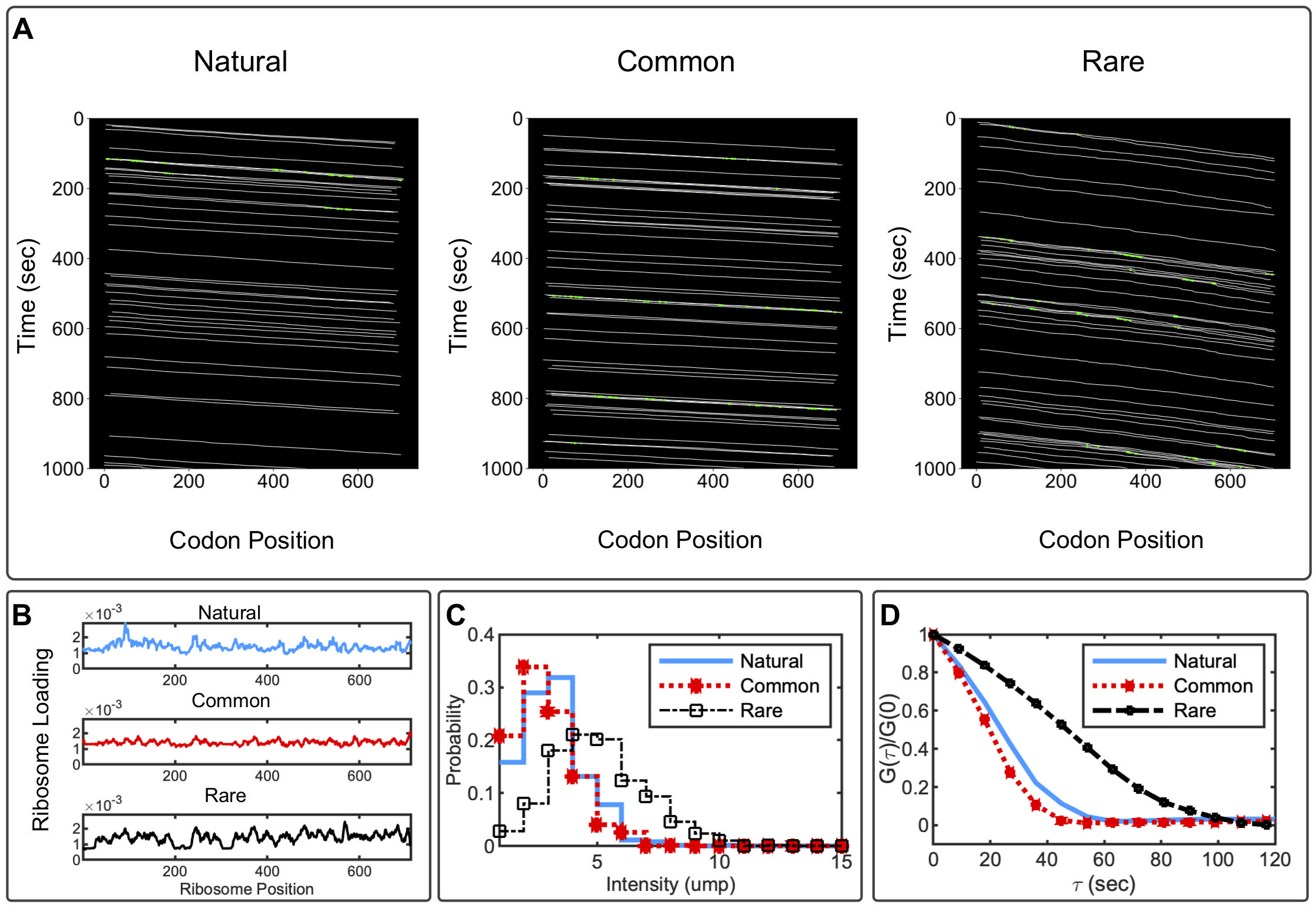

### Supplemental Fig S5

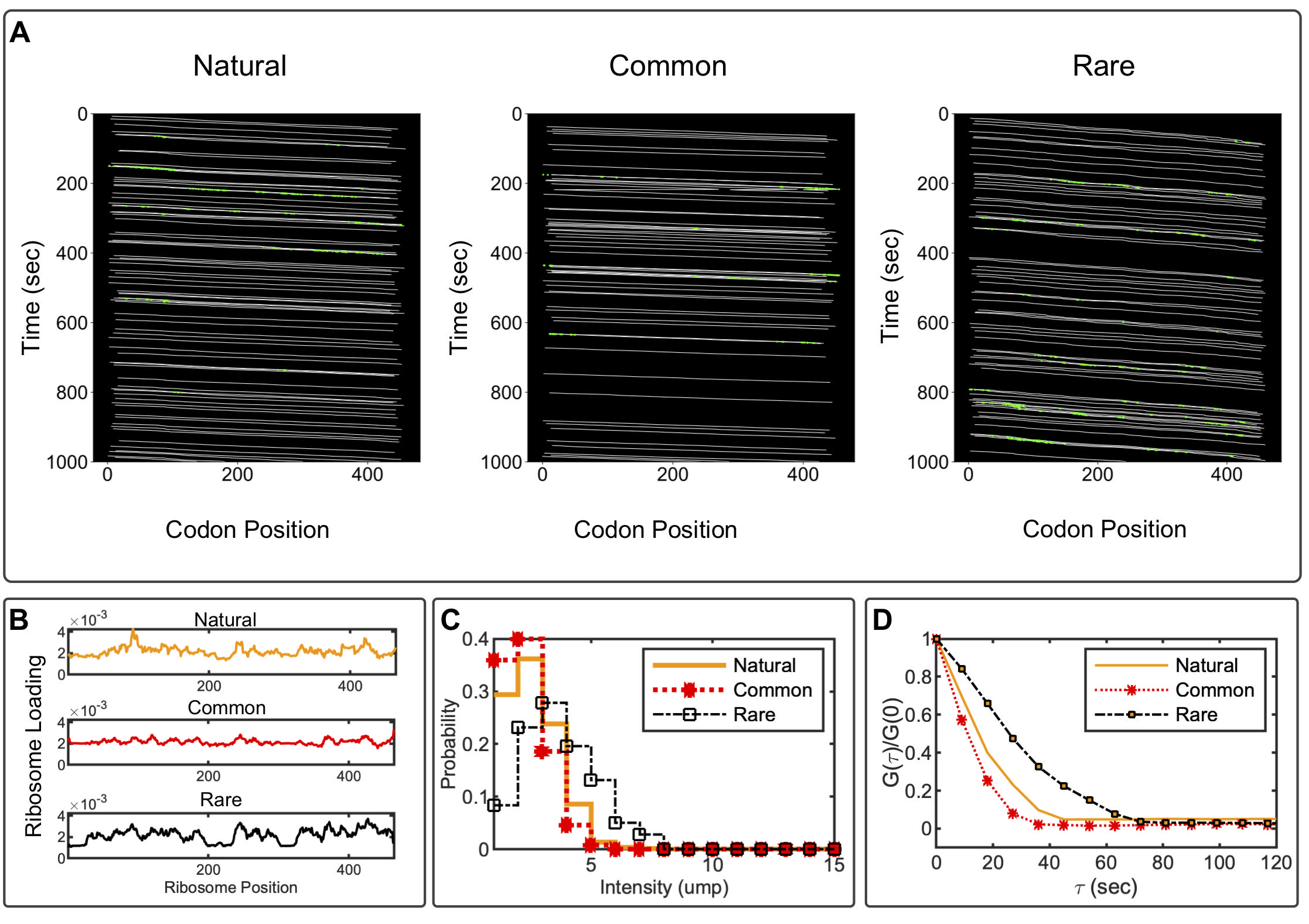

### Supplemental Fig S6

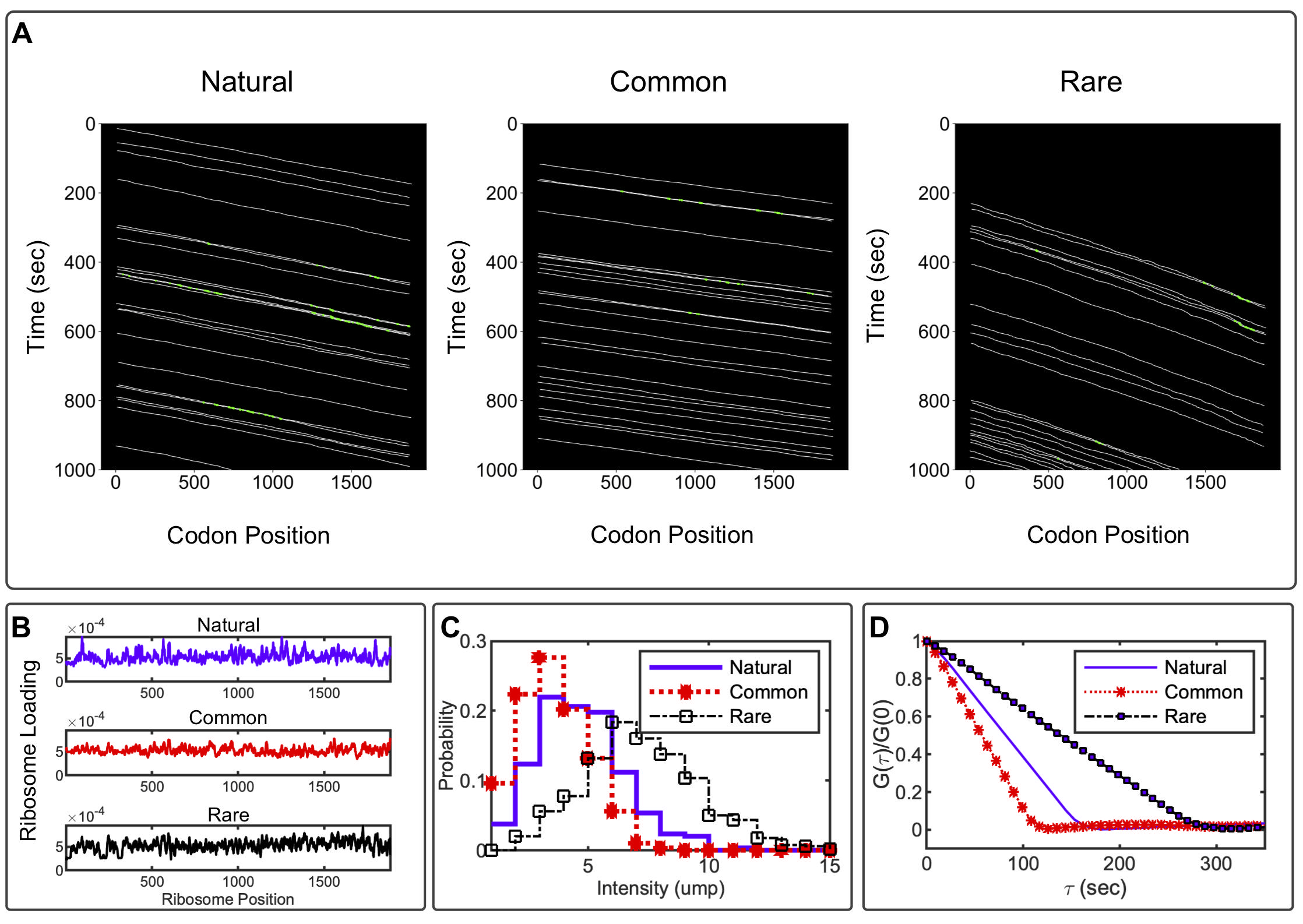

### Supplemental Fig S7

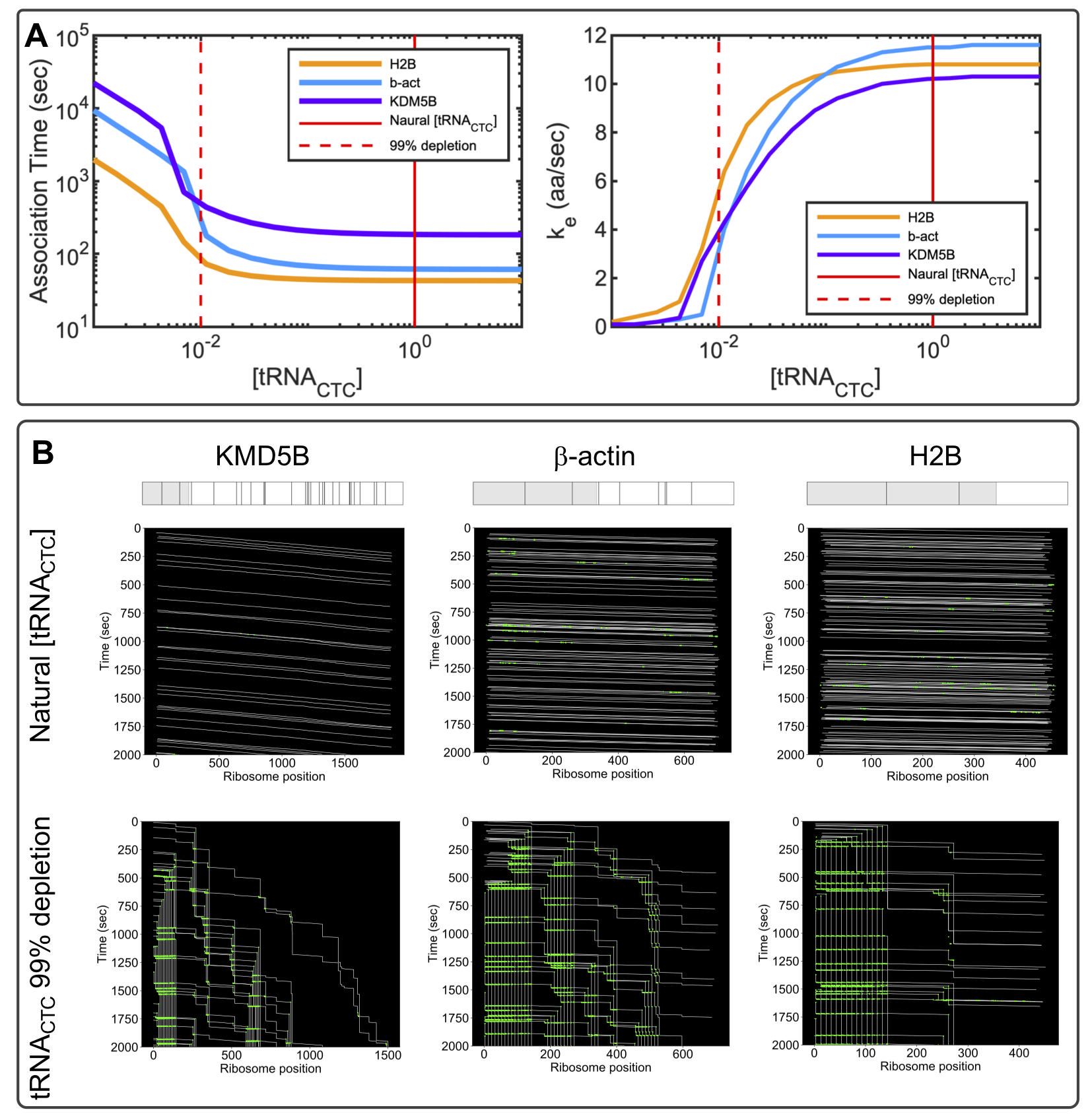

### Supplemental Fig S8

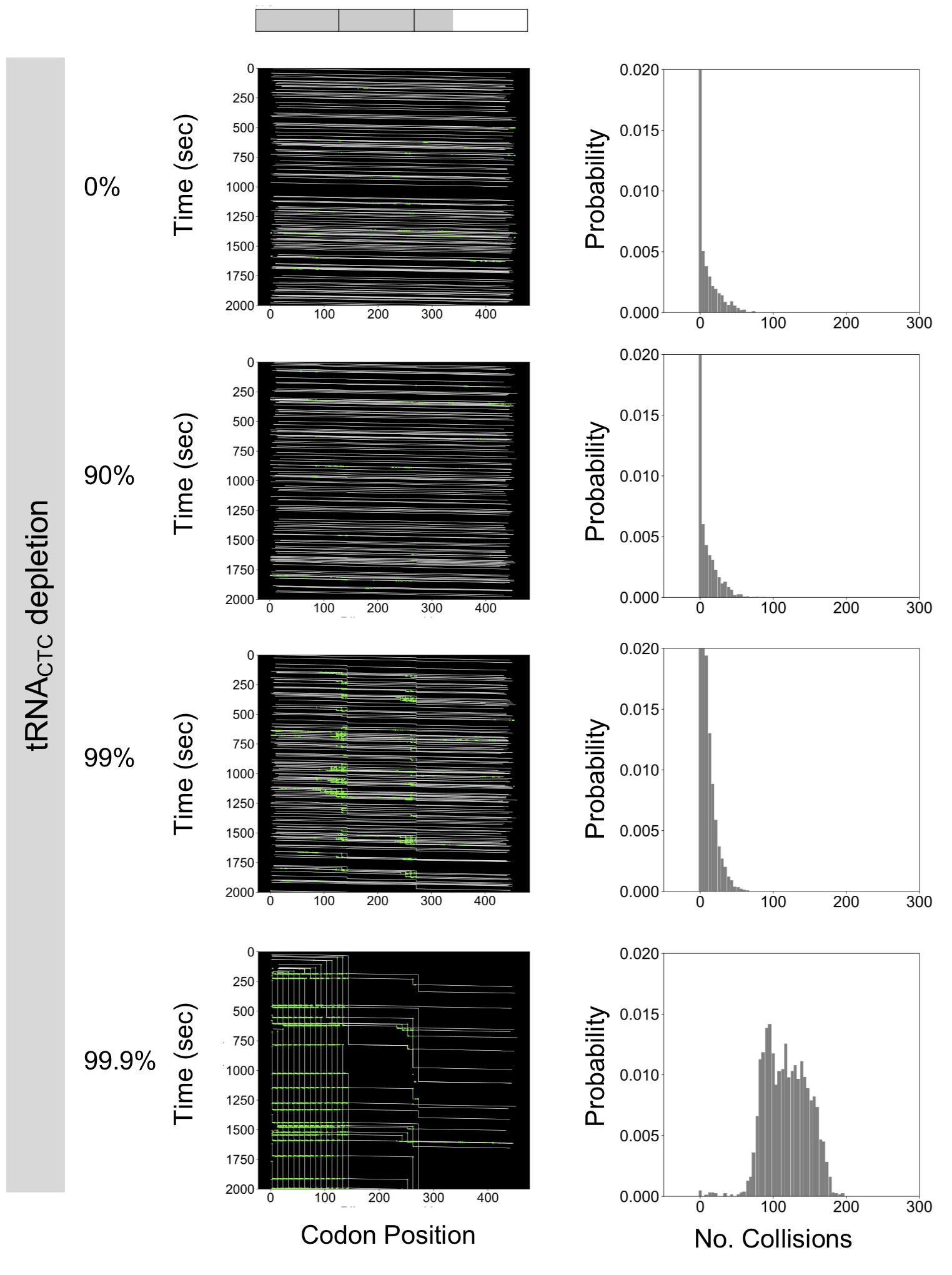

### Supplemental Fig S9

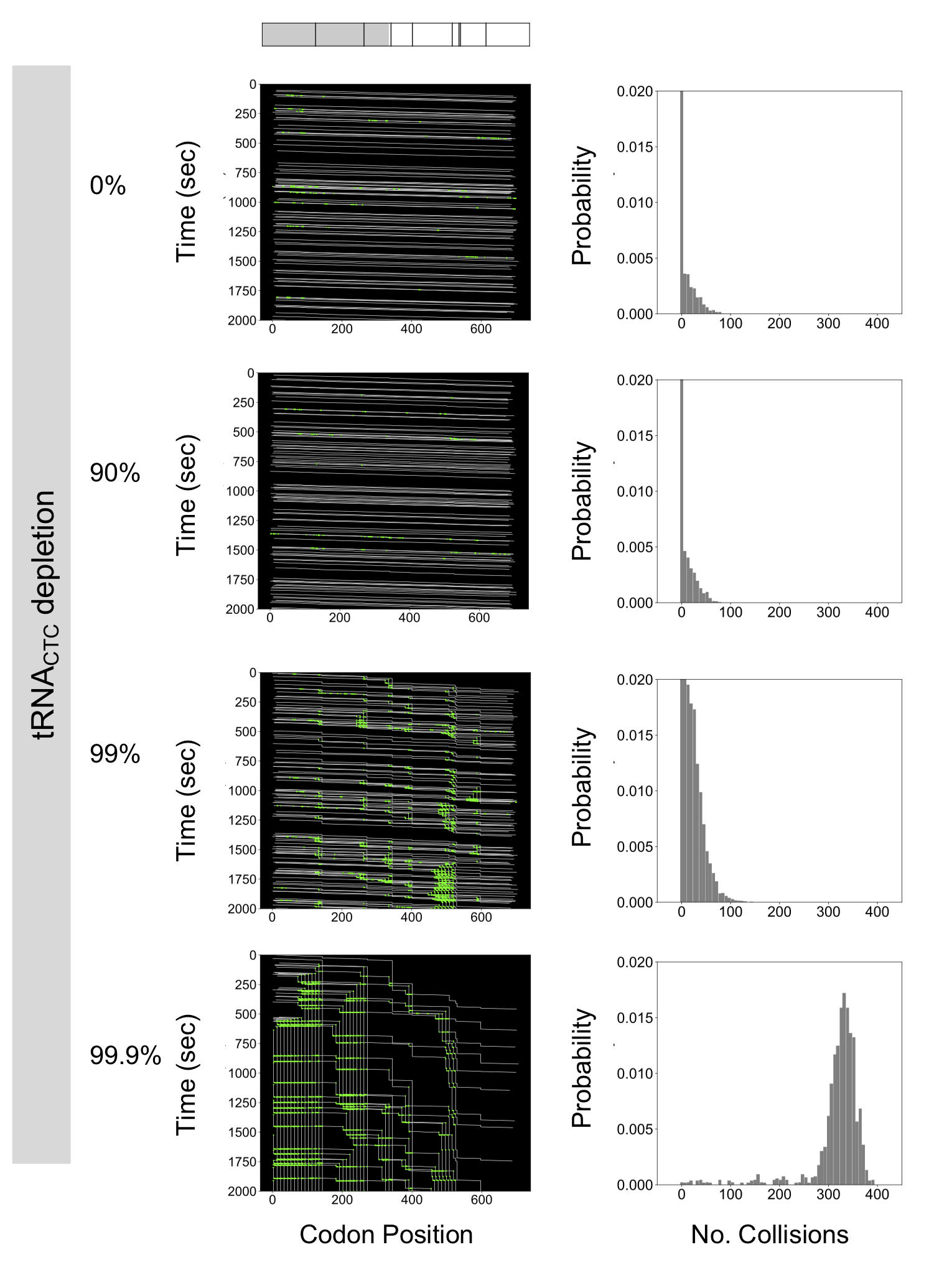

### Supplemental Fig S10

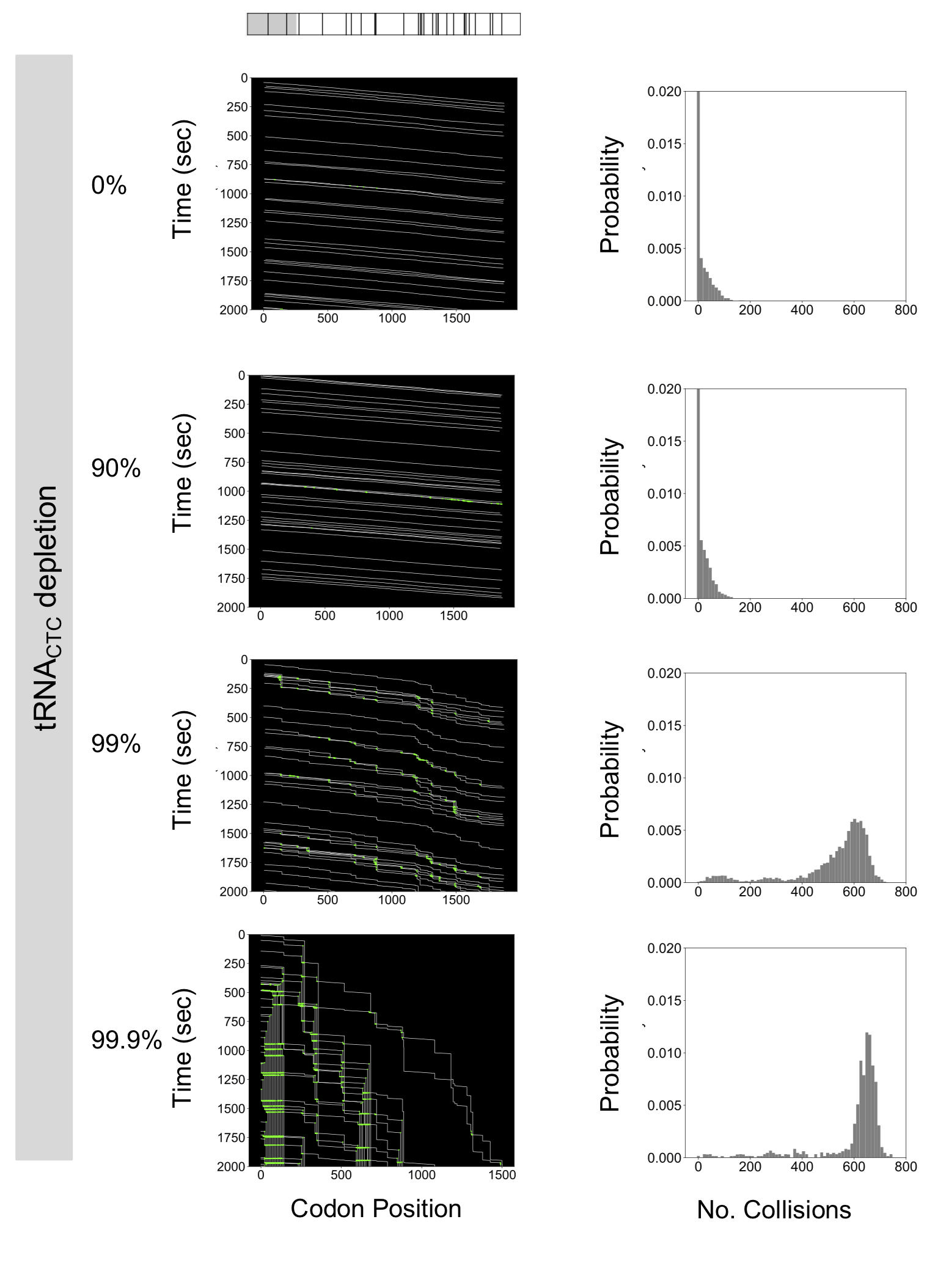

### Supplemental Fig S11

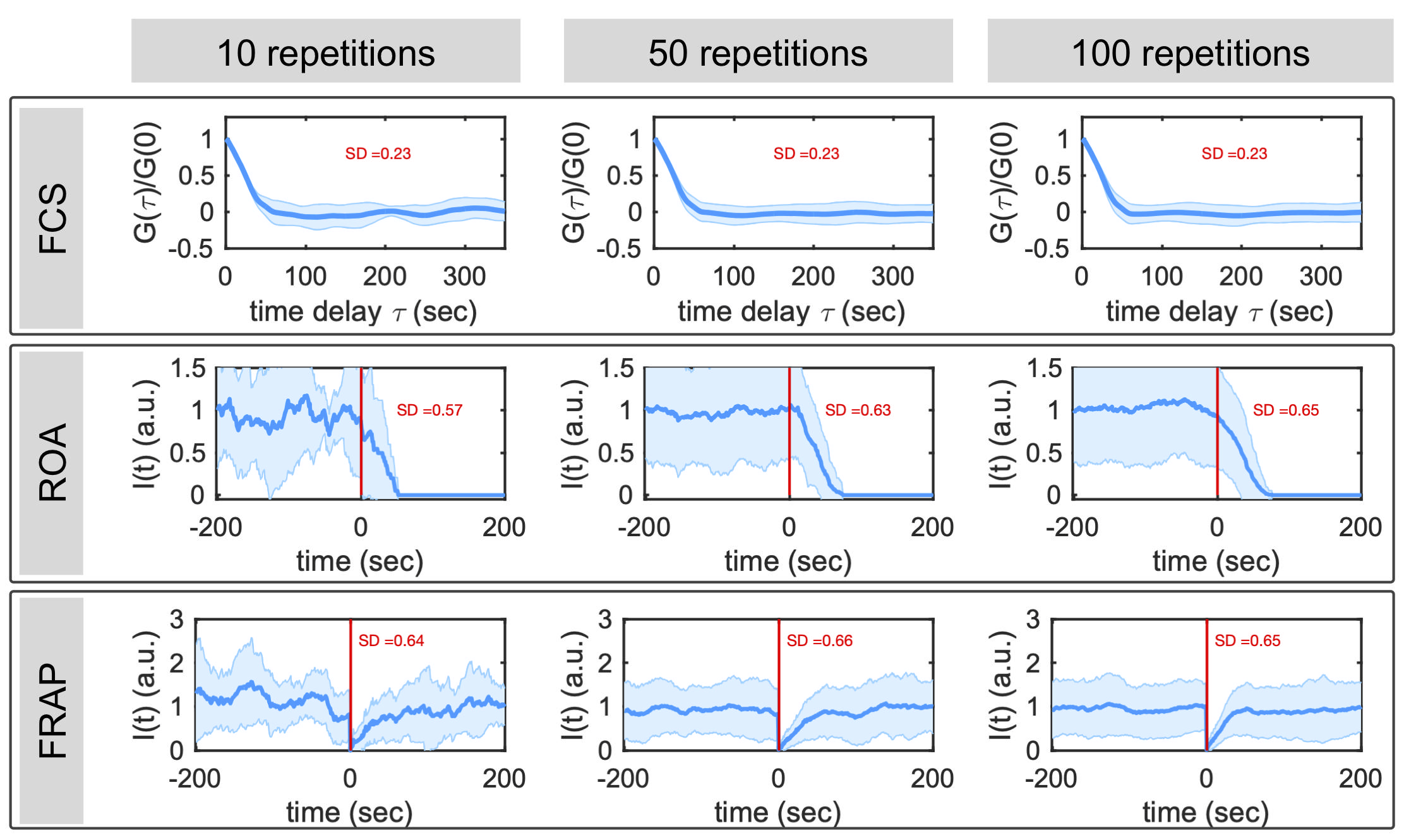

### Supplemental Figure S1

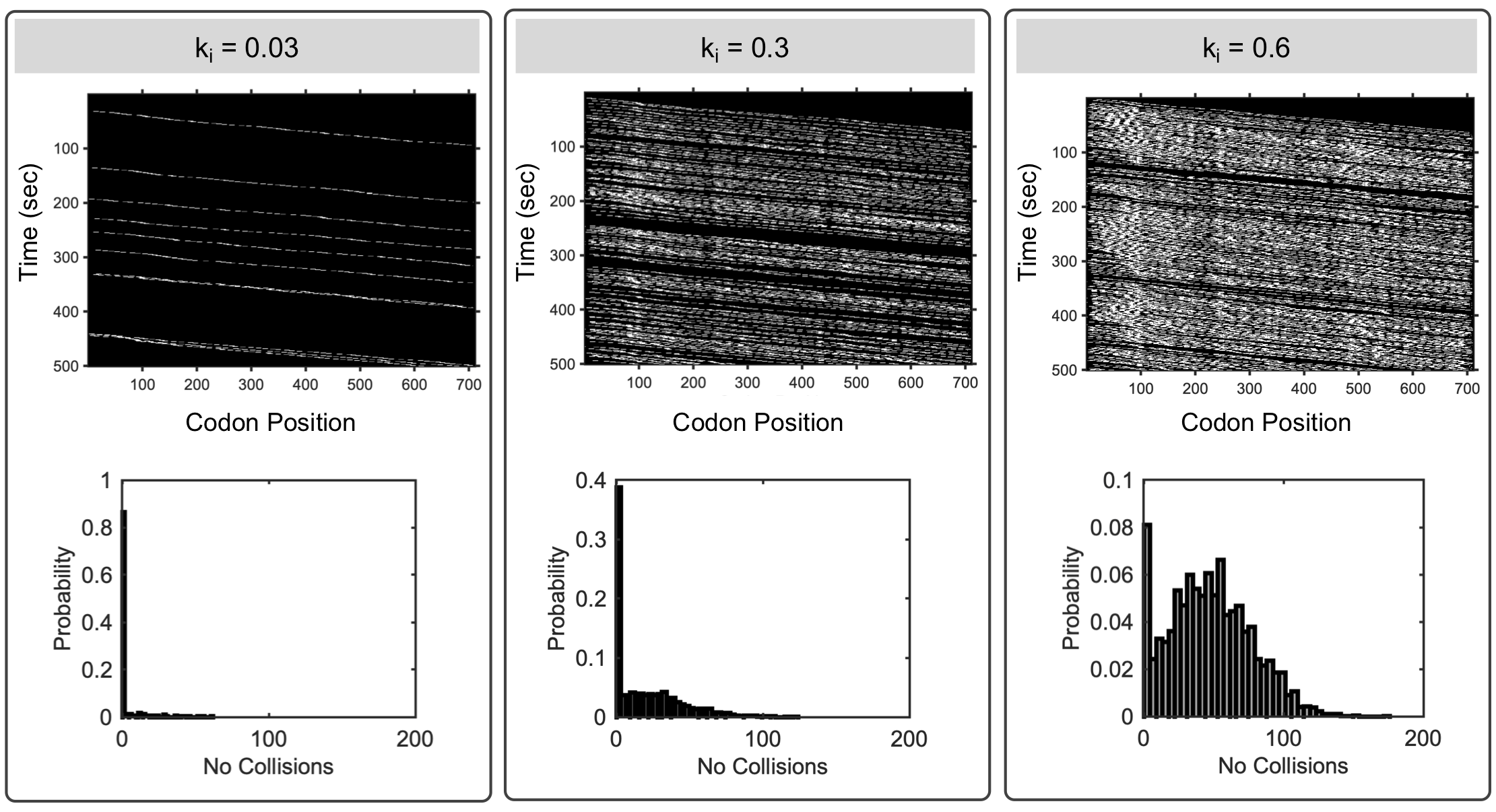

### Supplemental Figure S2

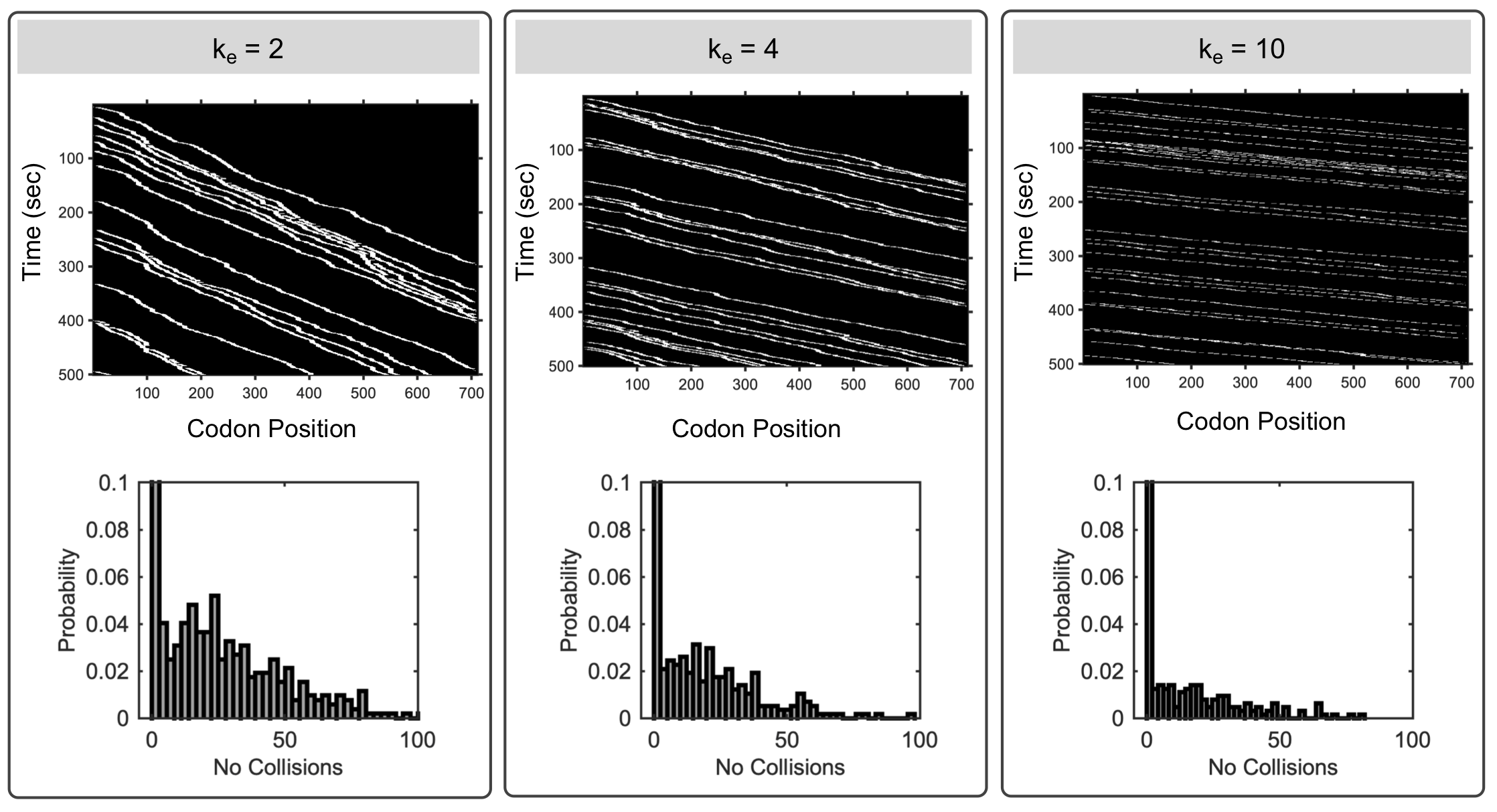
